# Development of the first antibody targeting vitellogenin using the *Palinurus elephas* mRNA molecular pathway

**DOI:** 10.1101/2021.08.26.457825

**Authors:** Faustina B Cannea, Cristina Follesa, Cristina Porcu, Rossano Rossino, Alessandra Olianas, Antonio Rescigno, Alessandra Padiglia

## Abstract

Vitellogenin is an essential protein involved in ovary maturation in many animals. Detection of this protein correlated with reproductive capacity may be important if carried out on marine organisms such as the red spiny lobster *Palinurus elephas*, a crustacean economically important crop from wild fish catches. Moreover, in recent years, vitellogenin has assumed an important role as a possible biomarker of marine environmental pollution, as its expression levels can be influenced by the presence of similar oestrogen pollutants and affect the reproductive sphere of marine organisms such as crustaceans. The *P. elephas* vitellogenin protein and its coding gene have never been isolated, so there is little information about its presence in this lobster. The aim of the present study was to develop a molecular strategy to create, for the first time, an antibody for the detection and quantization of vitellogenin in *P. elephas*.

## INTRODUCTION

The determination of vitellogenin (VTG) levels in marine aquatic species has for some time aroused much interest from both the fishery and environmental points of view. The study of VTG has now definitively found wide application in controlling the reproductive biology of fish, including the management of natural populations, the development of appropriate breeding practices and the quality control of the aquatic environment (Hiramatsu et al., 2002). VTG is a physiological indicator of the maturation stage of the female gonad (Reading et al., 2017) therefore, the possibility of measuring its concentration variations in many species throughout the life cycle makes VTG a far more sensitive indicator of sexual development with respect to endocrine parameters such as oestradiol, testosterone and gonadotropins (Denslow et al., 1999; Matozzo et al., 2008). VTG is a glycolipophosphoprotein (300-700 kDa) synthesized in the reproductive phase of many vertebrate and invertebrate animals. Expressed in the ovary and in the extraovarian sites of females in decapod crustaceans (Guan et al., 2016; Jia et al., 2016), VTG exhibits similar molecular features in phylogenetically related organisms, signifying its important physiological role in the course of evolution (Chen et al., 1997). Extraovarian synthesis involves the transport of VTG into the ovary through a blood vessel or haemolymph, in which it is internalized in the oocyte via receptor-mediated endocytosis (Barber et al., 1991; Ruan et al., 2020). Inside the oocyte, VTG undergoes a post-translational modification that consists of glycosylation events, the addition of lipids and proteolytic cleavages that generate smaller proteins collectively called vitelline (Vt), which are the main protein component for the development and maturation of eggs (Meusy et al., 1988; Wilder et al., 2002; Tiu et al., 2008). The proteolytic cuts originating from Vt occur in specific domains of the VTG precursor. The VTG amino acid sequence of the lobster *Homarus americanus* (HaVg1) deduced from cDNA highlighted the presence of several potential cleavage sites of the subtilisin endopeptidase (characterized by the consensus sequences RXXR or RXK/RK/R) from which at least three subunits with estimated molecular sizes of 80, 105 and 90 kDa could be produced starting from the precursor (Tiu et al., 2009). The precursor VTG is expressed at low levels in males and immature females, while it reaches high levels of expression during oogenesis in mature females (Thongda et al., 2015). Oestrogen 17 β-oestradiol and oestrogen-like molecules are powerful inducers of vitellogenesis in immature females since they can stimulate the expression of VTG (Palmer et al., 1998). In males, the genes for VTG (Vtgs) are normally silent, but in response to oestrogen or oestrogen-like substances, they undergo excessive transcriptional activity, and as a result, the protein synthesis occurs in the liver or functionally similar organs. Abnormal expression of this protein in males makes VTG a strong biomarker due to its ability to undergo variations in gene expression in aquatic species that live in aquatic environments contaminated by oestrogens and oestrogen-like molecules (Denslow et al., 1999; Sumpter and Jobling, 1995). In recent decades, greater attention has been given to assessing the adverse effects of chemicals that interfere with the endocrine system (EDCs: endocrine disrupting chemicals) in aquatic environments (Park et al., 2019). The concentration of VTG measured in the plasma of male fish under physiological conditions is 10-50 ng/mL, while in reproductive females, it is approximately 20 mg/mL (Denslow et al., 1999; Folmar et al., 1996; Parks et al., 1999). Studies on *Homarus americanus* have shown that VTG is undetectable in haemolymph from adult males instead increased 40-fold during the reproductive phase (Tsukimura et al., 2002). Since VTG levels have been shown to be closely related to the reproductive conditions of different crustaceans (Tsukimura, 2001; Tsukimura et al., 2002), their quantification could be a useful tool for monitoring the reproductive status of species living in specific environments. The European lobster *Palinurus elephas* (Fabricius, 1787) is a long-lived and slowgrowing species typical of temperate waters that is widely distributed in the Mediterranean Sea and the Atlantic Ocean. The development of increasingly powerful and efficient fishing methods in recent years has led to a decrement in the stocks of this crustacea in Sardinia (Italy) and other countries of the Mediterranean basin (Cau et al., 2019).

The main objective of this research was to develop a specific antibody for the detection of VTG in the eggs of the P. elephas lobster since the study of this protein has now definitively found wide application not only in controlling the reproductive biology of marine animals but also in monitoring the quality of aquatic environments. A specific antibody for *P. elephas* VTG could be a useful tool for the following reasons: 1) to investigate the reproductive suitability of the population of this crustacean since VTG is a good indicator of female reproductive activity, and 2) to evaluate nonphysiological protein amount in males since VTG is considered a biomarker of environment pollution. Moreover, the isolation of mRNA coding for VTG allowed us for the first time in this crustacean to enrich the gene databases with a part of the coding sequence from which it was possible to derive the corresponding primary structure of the encoded protein.

## MATERIALS AND METHODS

The VTG content present in the eggs of *P. elephas* was determined in ELISA experiments through the use of specific antibodies designed with a molecular strategy developed in our laboratory. Unlike standard methods that require the purification of proteins from specific tissues as a starting point for the production of specific antibodies, the method we used exploits synthetic peptides of which the primary sequence was deduced for the first time in our laboratory starting from a fragment of *P. elephas* Vtg gene (GenBank accession number: KX792013.1), which corresponds to the N-terminal region of others VTG sequences deposited in GenBank. Synthetic peptides have been used as antigenic molecules for the production of specific polyclonal antibodies.

### Animals

The study was conducted on the Mediterranean *P. elephas* egg clutches samples that were provided by the Marine Biology section of the Department of Life and Environmental Sciences of the University of Cagliari. The animals were captured in the period October to March, corresponding to the extruding period of the species (Goñi et al., 2003) inside and outside of two different fully protected areas (FPAs) located in the central western (Su Pallosu) and in the southwestern coasts (Buggerru) of Sardinia (Cau et al., 2019; Follesa et al., 2007). In particular, the concentration of VTG was estimated from the egg clutches of 30 ovigerous females at the same stage of intermediate development (stage 2) as follows: 16 ovigerous females were caught inside of FPAs (60.9.-91.9 mm carapace length) and 14 outside of FPAs (68.7-100.3 mm carapace length), all at the same developmental stage (Table S1A). The presence of external eggs on the pleopods and carapace length (CL) was used as an indicator of maturity (Kang et al., 2008). The developmental stage of the eggs of *P. elephas* (Follesa et al., 2007) was estimated using a binomial generalized linear model (R Core Team, 2017, https://www.R-project.org/) with a logistic link established through evaluating the von Bertalanffy growth parameters calculated for females. The relationship between the egg size at each developmental stage and the female lobster size was evaluated by regression analysis, and significant differences were calculated by t-tests (Zar, 1999). VTG levels were also determined in the hepatopancreas and haemolymph of 6 male *P. elephas*, of which 3 were captured inside and 3 were captured outside of the marine protected areas. Once collected, the biological samples were stored at -20°.

### Isolation of total RNA from ovarian tissue and RT-PCR

Total RNA was extracted from 50 mg of *P. elephas* eggs using TRI Reagent (Sigma-Aldrich, St. Louis, MO) following the manufacturer’s suggested protocol. The quality of purified RNA was verified by gel electrophoresis using a 1% denaturing agarose gel stained with SYBR Green II (Sigma-Aldrich), and the concentrations were measured using a NanoDrop 2000c UV-VIS Spectrophotometer (Thermo Scientific, Waltham, MA, USA) at 260 nm. To obtain cDNAs, *P. elephas* RNAs were reverse transcribed with an oligo dT primer using an enhanced avian myeloblastosis virus reverse transcriptase enzyme (Sigma-Aldrich) following the manufacturer’s instructions.

### *Palinurus elephas* Vtg cDNA amplification by PCR with hybrid primers

For detecting the unknown nucleotide sequences of the Vtg gene, a degenerate hybrid oligonucleotide primer (CODEHOP) strategy (Rose et al., 2003) was used, starting from aligning the multiple sequences of VTG proteins. In particular, nine different crustacean sources were chosen from the GenBank SwissProt database which were aligned using Clustal Omega (http://www.ebi.ac.uk/clustalw) (Fig. S1) and then cut into blocks using Block Marker software (http://blocks.fhcrc.org/blocks/). The primers were designed using the default parameters of the j-CODEHOP server (https://4virology.net/virology-ca-tools/j-codehop/). Amplification primers used for *P. elephas* Vtg cDNA were chosen from a group of primer candidates provided by the j-CODEHOP programme (Table 1).

**Table 1.**
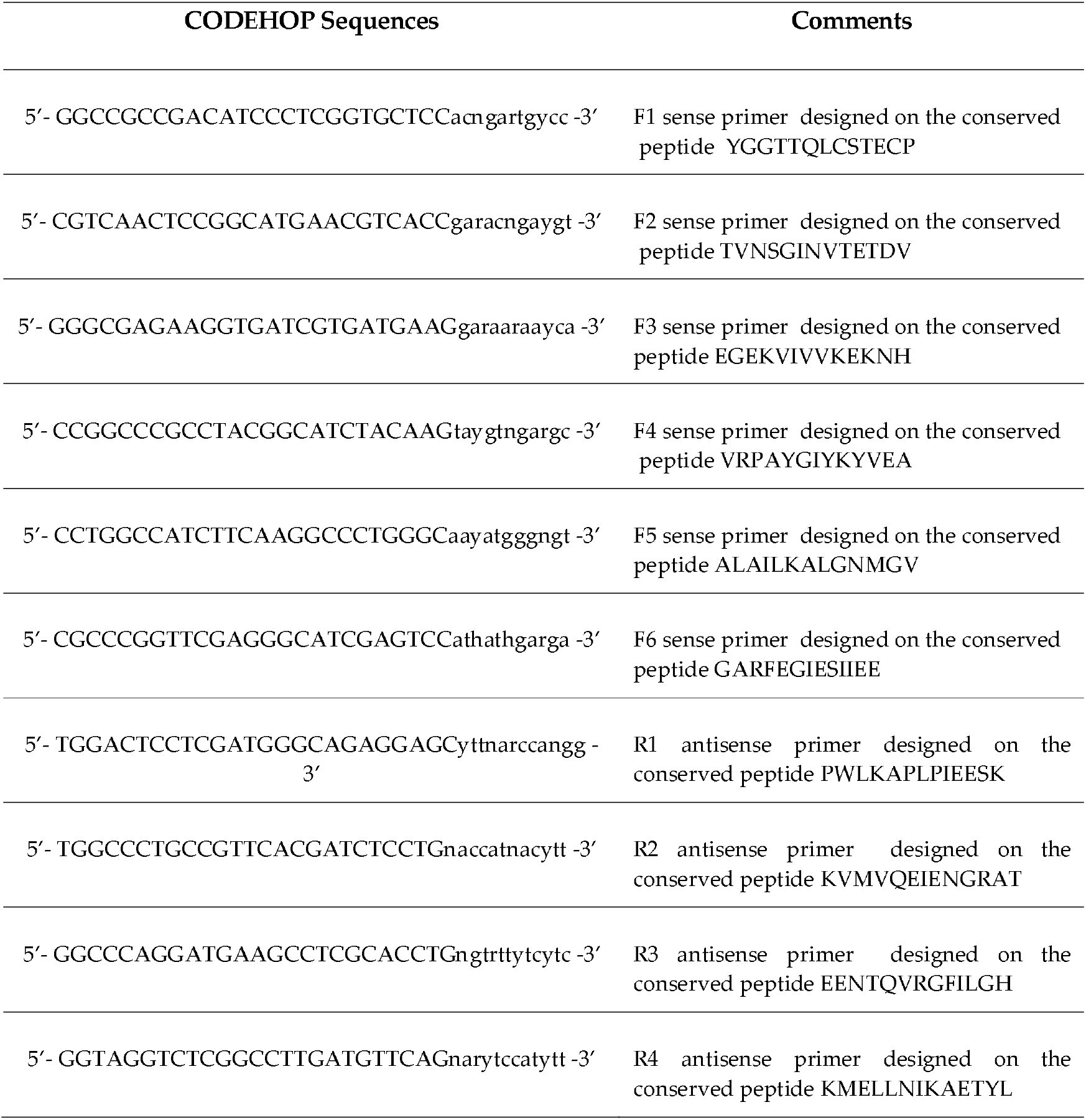
CODEHOP oligonucleotides used in PCR experiments and peptides chosen by the CODEHOP programme for the design of the primers. Each primer presents the consensus clamp given in the upper case, whereas the degenerate core is in the lower case: y = [C,T]; r = [A,G], and n = [A,G,C,T]. The position and orientation of each primer is reported in the Supplementary information (Fig. S1).

Each primer presents the consensus clamp given in the upper case, whereas the degenerate core is in the lower case: y = [C,T]; r = [A,G], and n = [A,G,C,T]. PCR was performed in a solution containing 1.5 mM MgCl_2_, 100 mM Tris-HCl, pH 8.3, 50 mM KCl, 200 mM dNTP mix, 1 mM sense primer, 1 mM antisense primer, 1 μg of *P. elephas* cDNA, and 1-3 units of Jump Start AccuTaq LA DNA polymerase mix (Sigma-Aldrich). Thermal cycles of amplification were carried out in a Personal Eppendorf Mastercycler (Eppendorf, Hamburg, Germany) using slightly different programmes. The PCR experiments, conducted using all the pairs of sense and antisense oligonucleotides chosen by j-CODEHOP software, allowed us to obtain a single reaction product with the F3-R2 primer pair (Fig. S1), with dimensions of approximately 300 bp, which was compatible with those of a possible expected product. The PCR products detected on 6% polyacrylamide or 2% agarose gels were purified with a Charge Switch PCR Clean-Up kit (Invitrogen, Carlsbad, CA) and then sent to BMR genomics (Padova, Italy) for sequencing. Translation of nucleotide sequences was performed using OMIGA or ExPASy translate routine software (http://ca.expasy.org/) (Fig. 1A). Sequences were aligned with Clustal Omega (https://www.ebi.ac.uk/Tools/msa/clustalo/), and similarities were analysed with the advanced BLAST algorithm available at the National Center for Biotechnology Information website (http://www.ncbi.nlm.nih.gov/) and with the FASTA algorithm v.3.0 from the European Bioinformatics Institute website (http://www.ebi.ac.uk/fasta33/index.htlm).

**Fig. 1.**
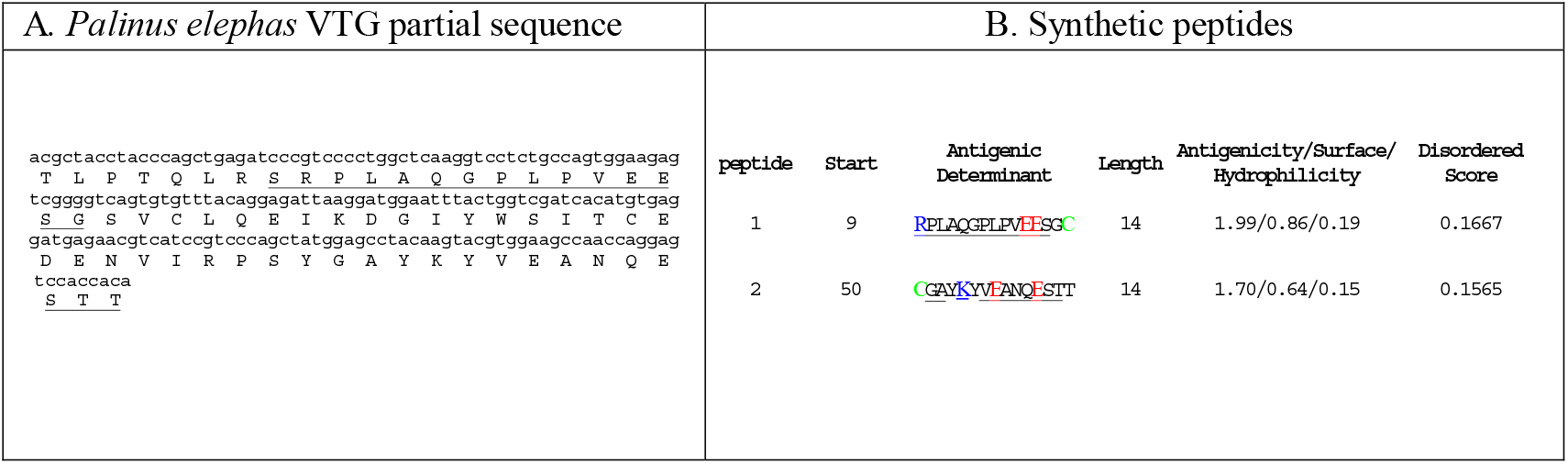
**(A) Nucleotides and, below, deduced amino acid sequences of VTG from *P. elephas***. The underlined regions are predicted to be antigenic regions. **(B) Information related to the synthetic peptides**. An extra ‘C’ (highlighted in green) is added to the C terminus (or N terminus) to facilitate conjugation. The antigenic positively charged residues (K, R) and negatively charged residues (E) are in blue and in red respectively. The synthesis and epitopes predictions were made using GenScript. Residues of VTG enzyme and synthetic peptides are reported with the one letter code.

### Peptide synthesis

The *P. elephas* VTG amino acid sequence deduced in silico (Fig. 1A) was used to create two synthetic peptides (Fig 1B) (Genscript USA Inc.) to be employed for the production of specific anti-VTG antibodies (Twin Helix, Rho, Italy). The epitopes were predicted by the GenScript Optimum AntigenTM design tool (antigenic peptide 1 AA9-AA23 and antigenic peptide 2 AA50-AA63). Comprehensive analysis was performed on multiple aspects, including antigenicity, hydrophilicity, hydrophobicity, the probability of antibody accessibility (exposure on the protein surface) and the uniqueness of the protein sequence. The useful information related to the synthetic peptides for the synthesis of antibodies is shown in Fig. 1B.

### *P. elephas* VTG detection by ELISA

For the detection of VTG, we performed indirect ELISA using the commercial Prepro Tech TMB ELISA Buffer Kit (DBA, Italy) following the manufacturer’s suggested protocol. For the standard curve, the antigenic peptides diluted in 2% blocking medium in phosphate buffered saline buffer (PBS, pH 7.4) were added to the wells of the plate at scalar concentrations in the range of 1-250 ng per well and then incubated for one hour at 37 °C. After three consecutive washes in PBS, the antibodies diluted in 2% blocking medium in PBS at a concentration of 0.1 µg/well were incubated on the plates overnight at 4 °C. After the incubation period, the plates were washed five times, and secondary antibodies made of anti-mouse IgG conjugated with HRP diluted in PBS (1:20000) were added to the wells of the plate for 2 hours at 37 °C. Finally, the plates were washed six times and then incubated with the tetramethylbenzidine (TMB) substrate HRP. The reaction was carried out in phosphate-citrate buffer (pH 5.5) in the dark for 15 minutes at room temperature and subsequently blocked with 10% sulfuric acid. The plates were read at a wavelength of 460 nm using the VICTOR 3 V 1420 Multi-Label Microplate Reader (Perkin Elmer, USA). All reactions were carried out in triplicate using both antigenic peptides in separate experiments. The blank samples were composed of buffer or water with no protein sample included. The standard curve R2 values obtained were the same for antigenic peptide 1 (R2= 0.9757) and peptide 2 (R2= 0,97). When the homogenate of *P. elephas* eggs was used as an antigen, the ELISA plates were coated with 25 mg of biological sample per well. Specifically, the eggs were sonicated in RIPA buffer supplemented with a protease inhibitor (Bio-Rad Laboratories, Inc.) in the proportion of 25 mg of eggs per 100 microlitres of buffer. After homogenization, the tubes were centrifuged at 14000 x g for 20 minutes, and the supernatant obtained was used following the steps previously described for the peptides used to construct the standard curve. To validate the ELISA, we determined the intra-assay coefficient of variation (CV) value, which was less than 15%.

## Results AND DISCUSSION

The aim of the present work was to design a specific antibody for the detection of VTG in *P. elephas* females at the same stage of intermediate development (stage 2). The samples with external eggs in the intermediate stage (diameter 0.92-1.30 mm) were selected since the levels of VTG in crustaceans appear to decrease significantly with the beginning of the oviposition of the animals (stage 3) (Tsukimura et al., 2002). The objective was achieved after isolating the coding mRNA, which, after being reverse transcribed, was subjected to PCR that utilized copies of partially degenerated primers designed with the CODEHOP strategy (Rose et al., 2003) starting from aligning the multiple sequences of VTG proteins (Fig. S1). The results of the PCR fragment sequencing allowed us to reconstruct in silico a part of the primary structure of the protein, corresponding to a peptide of 63 amino acids (Fig. 1A). By aligning the sequence obtained with protein sequences deposited in NCBI databases (https://blast.ncbi.nlm.nih.gov/Blast.cgi) and UniProt (https://www.UniProt.org/), we found a homology identity of 60-70% with the amino terminal portion of other crustacean VTGs (Fig. S2).

Analysis of the amino acid sequence to establish the presence of antigenic regions revealed the potential presence of two epitopes. This allowed the biotechnological synthesis of two peptides (the antigenic peptide 1 AA 9-AA23 and antigenic peptide 2 AA50-AA63) (Fig. 1B) that were used to create two different antibodies (anti-VTG1 and anti-VTG2) for ELISA experiments to detect the VTG expressed in the females of *P. elephas* in the second stage of egg maturation. Different concentrations of peptide 1 or peptide 2 were used in the presence of the specific antibody (anti-VTG1 or anti-VTG2), to construct the standard reference curve. The standard curve R2 values obtained were the same for antigenic peptide 1 (R2= 0.9757) and peptide 2 (R2= 0,97). Biological samples used in ELISA experiments were obtained from 30 females and 6 males of *P. elephas* captured inside and outside of the marine protected areas surrounding the coasts of Sardinia. The external eggs attached to the pleopods were removed from the females; the haemolymph was taken from the males. By relating the absorbance of each sample with an unknown concentration to the standard curve previously established, we obtained the amounts (ng/mL) of VTG contained in the eggs of *P. elephas*. These were found in a range of 200-260 ng in 14 of the 30 females analysed, while in the other 16, the concentrations ranged from 120-180 ng/mL (Table S1 B). Differences in VTG levels in stage 2 eggs could be correlated with the sexual maturity of females within the same stage of development. The VTG concentration did not vary between the females captured inside (IN) and outside (OUT) FPAs. In fact, analysis of the VTG concentration in relationship with the inside and outside FPAs did not show statistic differences (Fisher’s exact test statistic = 1; p > 0.05). According to those found for Homarus americanus males (Tsukimura et al., 2002), the absorbance values obtained for *P. elephas* males were so low that it was not possible to calculate concentration values to correlate with the standard curve. All ELISA experiments were conducted in parallel using both available antibodies, and the VTG concentrations detected were the same in the presence of each anti-VTG antibody. An overview of steps involved in the development of *P*.*elephas* anti-VTG antibodies is shown in Fig. 2.

**Fig. 2.**
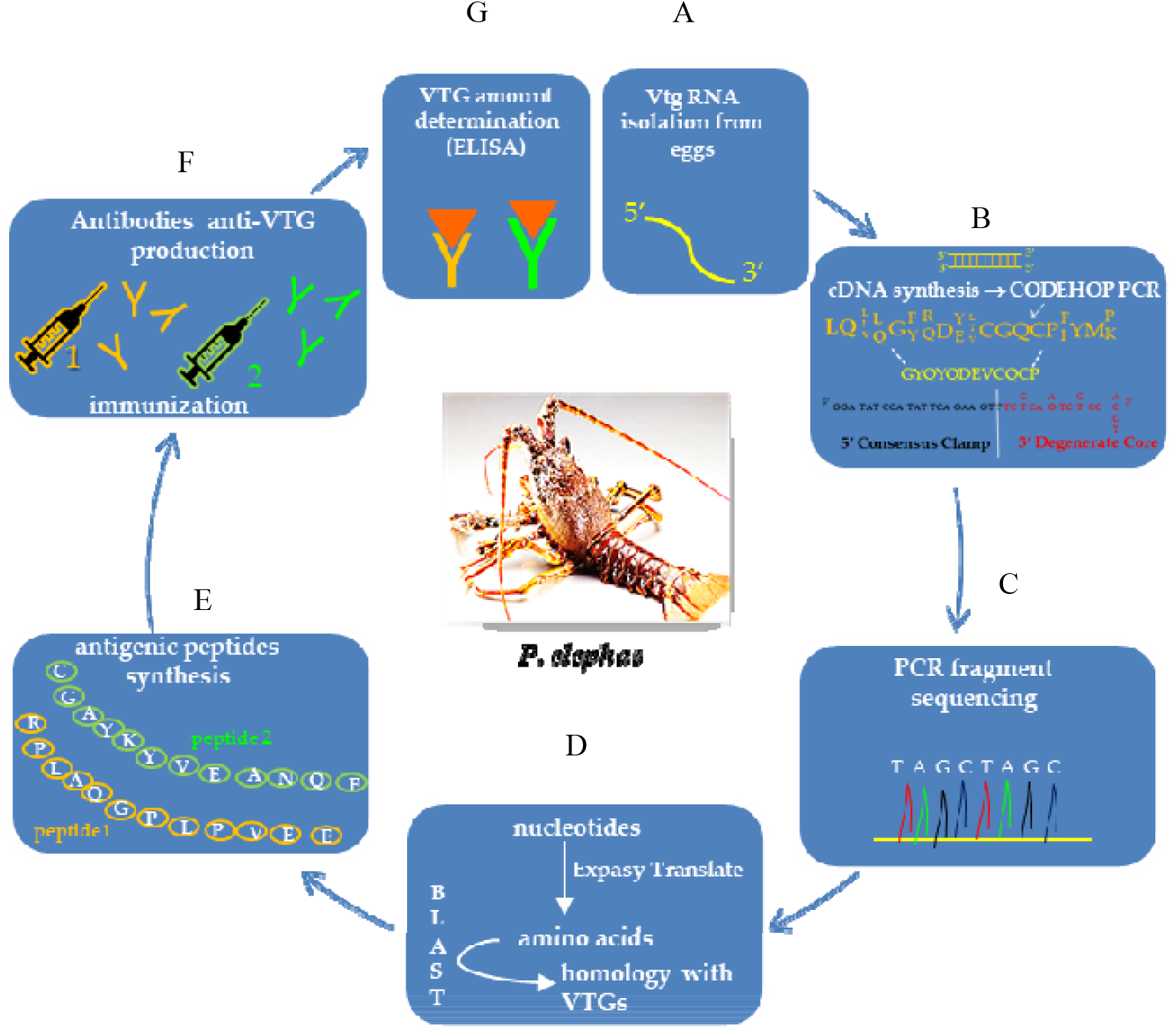
An overview of steps involved in the development of *P*.*elephas* anti-VTG antibodies. (A) Total RNA was extracted from *P. elephas* eggs. (B) To obtain cDNAs, *P. elephas* RNAs were reverse transcribed; for detecting the unknown nucleotide sequences of the Vtg gene, CODEHOP PCR was used, starting from aligning the multiple sequences of VTG proteins. (C) The PCR products obtained were used for the preparation of sequencing samples. (D) The nucleotide sequence acquired experimentally was translated into the amino acid sequence and aligned with protein sequences deposited in NCBI databases. (E) The potential presence of two epitopes in the virtual VTG peptide allowed the biotechnological synthesis of two peptides (peptide 1 and peptide 2) which were used to create two different antibodies for ELISA experiments (F) to detect the VTG in the females of *P. elephas* in the second stage of egg maturation.

## Conclusion

In this work, a study aimed at the creation of the anti-VTG antibodies of *P. elephas* was carried out for the first time starting from the partial isolation of the mRNA coding for VTG. This strategy allowed a part of the primary structure of the protein to be defined and the biotechnological synthesis of two peptides that, thanks to their high antigenic potential, were used to generate two different antibodies. With this molecular pathway, we have bypassed the protein purification procedure from tissue or other biological samples where the protein is expressed, which is usually used to generate antibodies (Denslow et al., 1999; Tsukimura et al., 2002; Kang et al., 2008). As reported in the literature (Denslow et al., 1999), the quantization of VTG through ELISA represents a powerful tool that can be used to achieve further objectives in ecological and marine biological studies of *P. elephas*, due to the following:

1) The tool makes it possible to compare the levels seen in individuals of *P. elephas* taken from different repopulation areas, both in protected areas (internal) and in areas dedicated to commercial fishing (external), to verify the possible role of the protein in reproduction and growth and increase the number of individuals in the population itself.

2) The tool can monitor the concentrations of this protein in individuals grown in both restocking and free areas to verify whether any increase in protein levels could be related to the presence of oestrogen-like substances in the water.

In fact, the use of VTG as a biomarker can indicate not only the presence of a pollutant in the water but also the effect that it can have on physiological alterations related to the reproductive sphere. The VTG values obtained in the various samples during this study did not highlight any significant differences between the expression levels of mature females living inside and outside protected marine areas. The values found allowed to establish, for the first time, a concentration range for VTG considering the eggs of females in ovarian stage 2, probably correlated with sexual maturity within the same stage of development. The analyses carried out on male individuals did not provide detectable results. A future goal will be to analyse a greater number of individuals to constantly monitor any fluctuations in the concentration of this important protein both in mature females and in males and to determine if environmental conditions determine its expression.

## Competing interests

The authors declare that they have no competing interests.

## Funding

This research was partially supported by FIR (Fondi integrativi per la Ricerca) funded by the University of Cagliari (Italy)

## Author Contributions

Conceptualization, F.B.C., A.P., C.F. and C.P.; methodology, F.B.C. and A.P.; software, R.R. and A.O; validation, F.B.C., A.P. and C.F.; formal analysis, F.B.C. and C.F.; investigation, F.B.C., A.P., C.F. and C.P; resources, C.F., A.P., R.R; data curation, F.B.C. and A.P.; writing—original draft preparation, AP.; writing—review and editing, A.R.; visualization, F.B.C., A.P. and C.F.; supervision, A.P.; project administration, A.P and C.F; All authors have read and agreed to the published version of the manuscript.

## Supplementary information

**Fig. S1.**
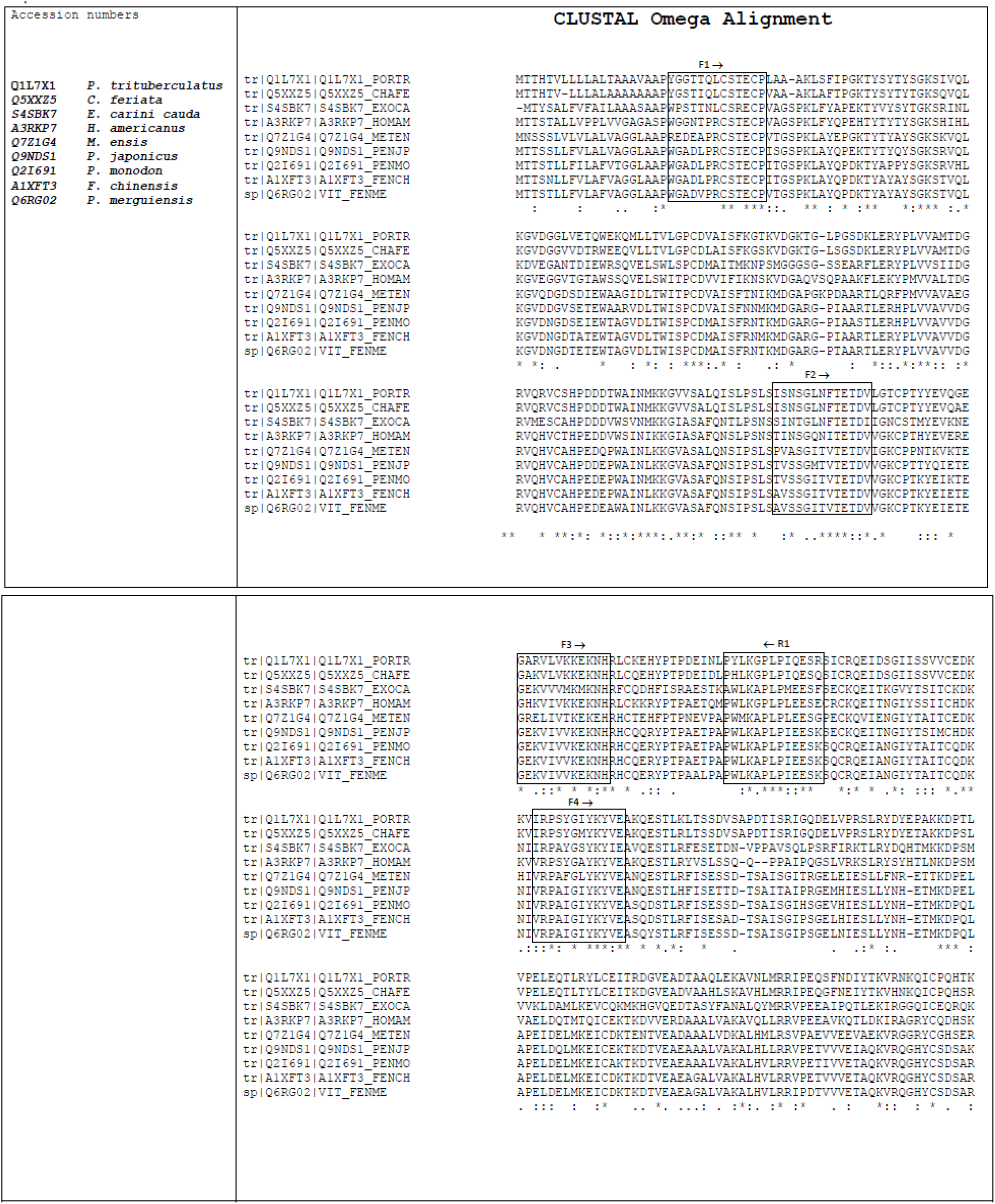
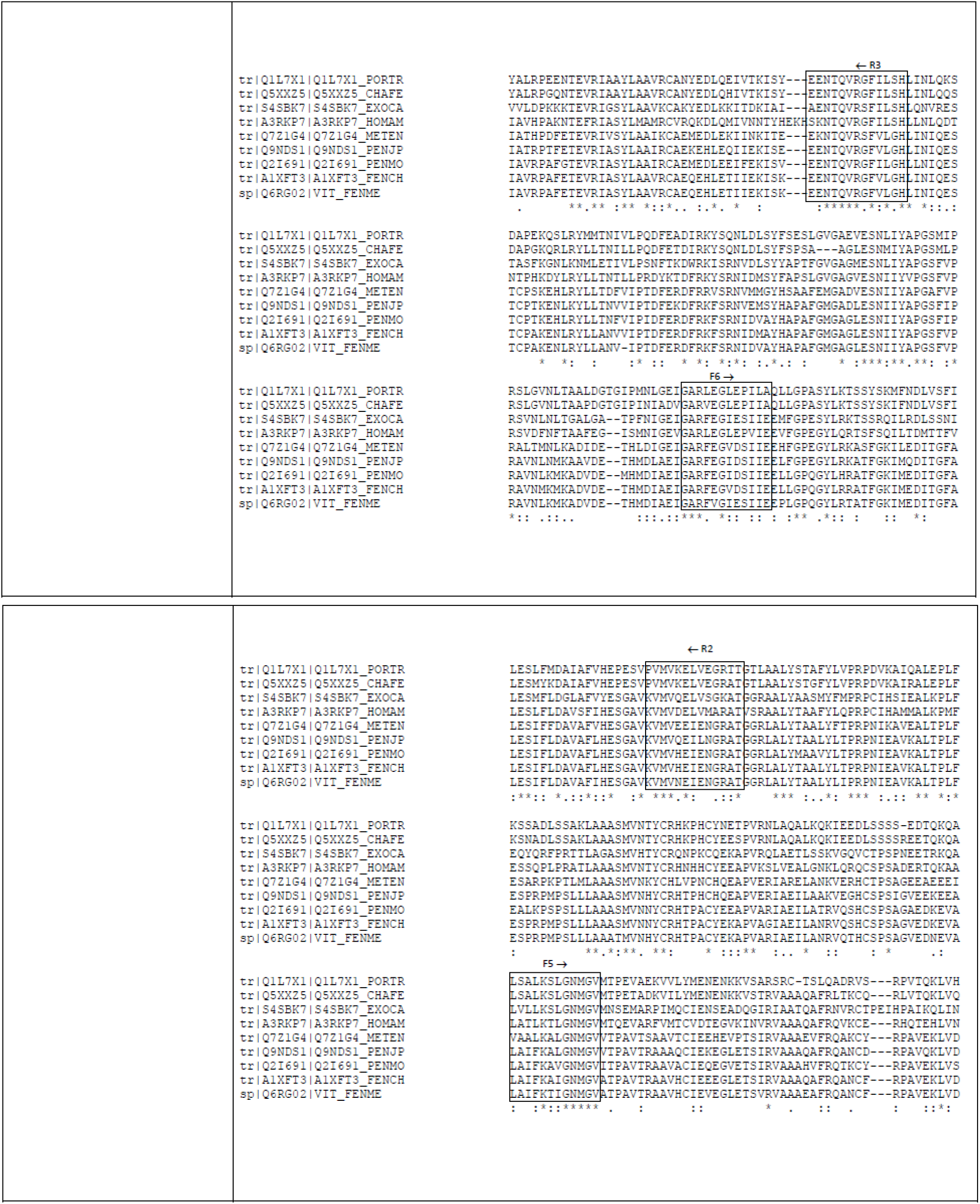
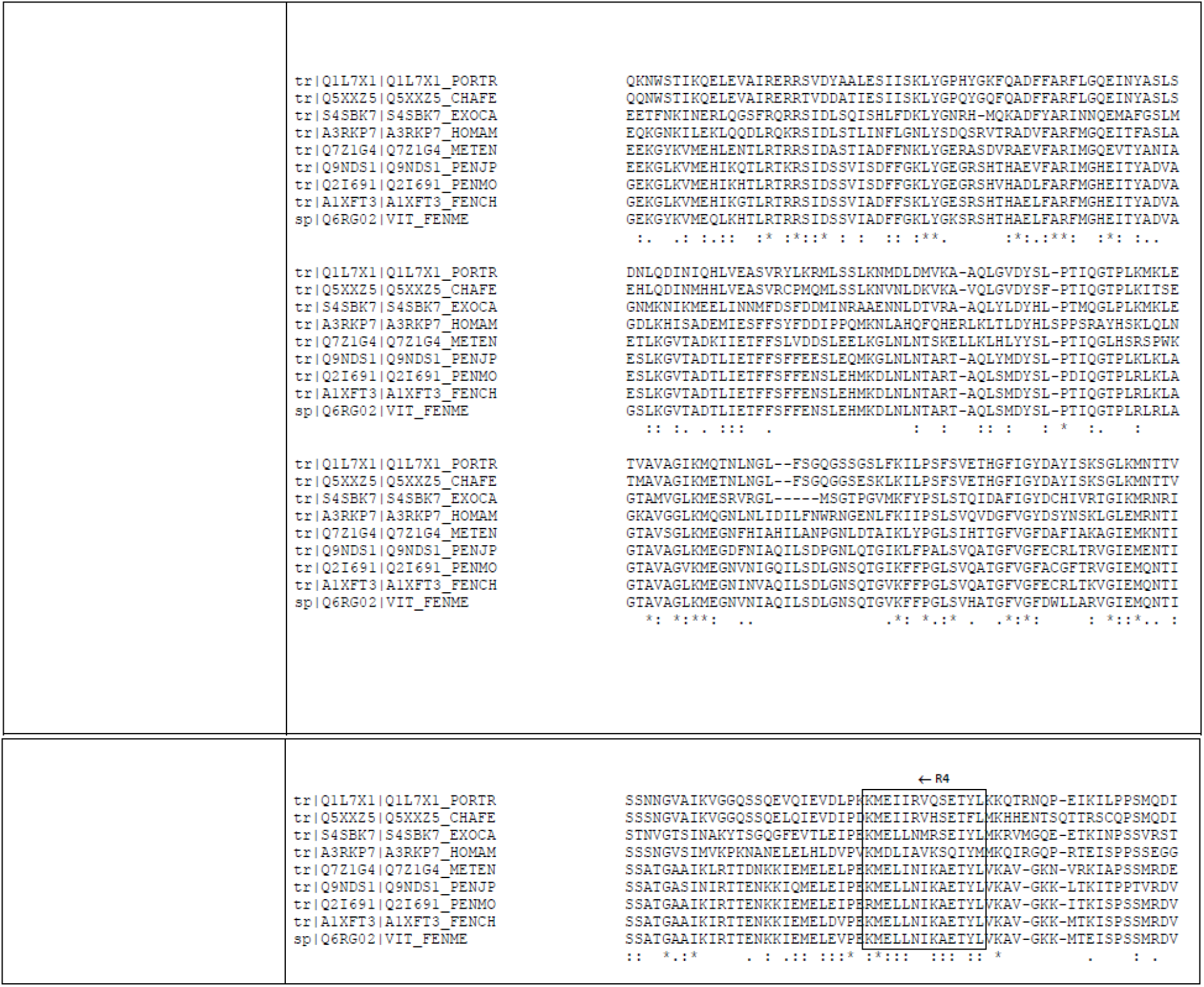
Multiple alignment of VTG amino acid sequences from nine different crustacean sources. The sequences were chosen from the GenBank SwissProt database and aligned using Clustal Omega (https://www.ebi.ac.uk/Tools/msa/clustalo/). The figure shows the alignments based upon which the CODEHOP primers have been designed. An asterisk (*) denotes identical residues; a colon (:) denotes a conserved residue substitution; a full stop (.) denotes partial conservation of the residue. The arrows above the amino acid sequences indicate the position of the sense (F →) and antisense (← R) primers chosen from a group of candidate primers obtained from the j-CODEHOP programme. The sequences of the sense and antisense primers, shown inside the boxes, were obtained from the Clustal Omega alignments.

**Fig. S2.**
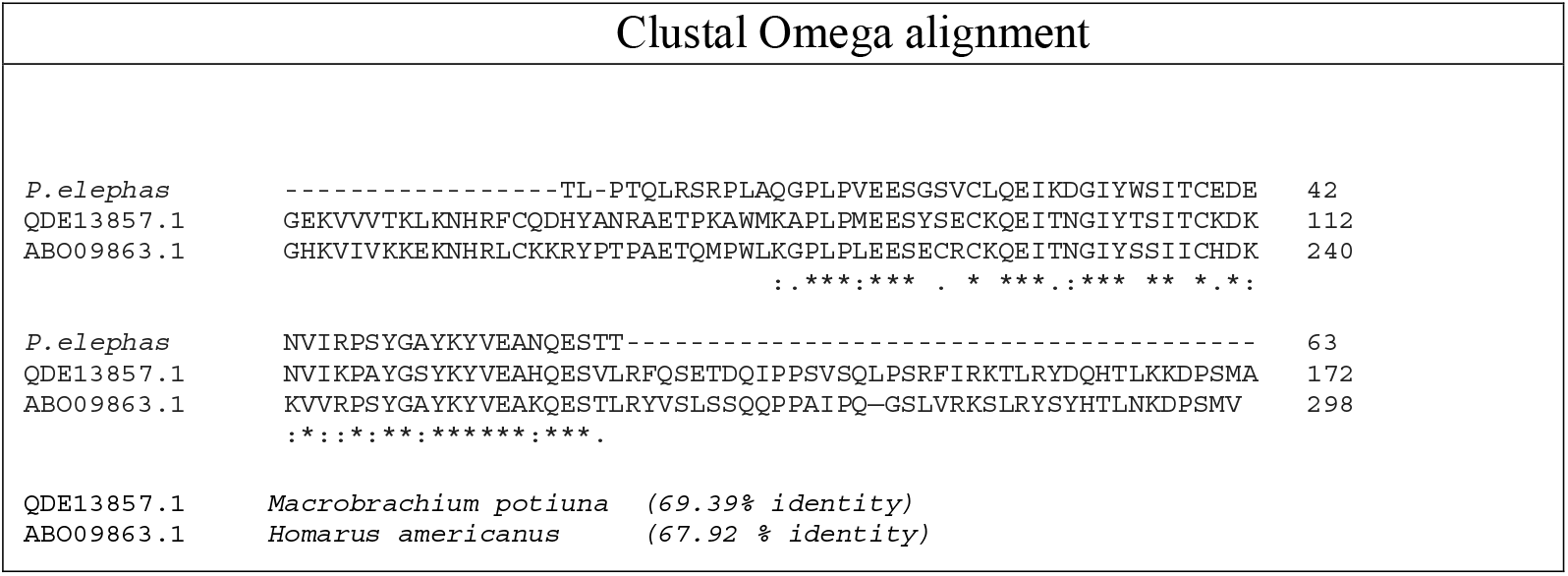
Alignment of VTG *P. elephas* amino acid sequences with homologous VTG sequences from GenBank. Identity and similarity percentages are shown at the top of each Clustal Omega alignment panel. Numbers on the left indicate the positions of the amino acids in each protein. An asterisk (*) denotes identical residues; a colon (:) denotes a conserved residue substitution; a full stop (.) denotes partial conservation of the residue.

**Table 1S.**
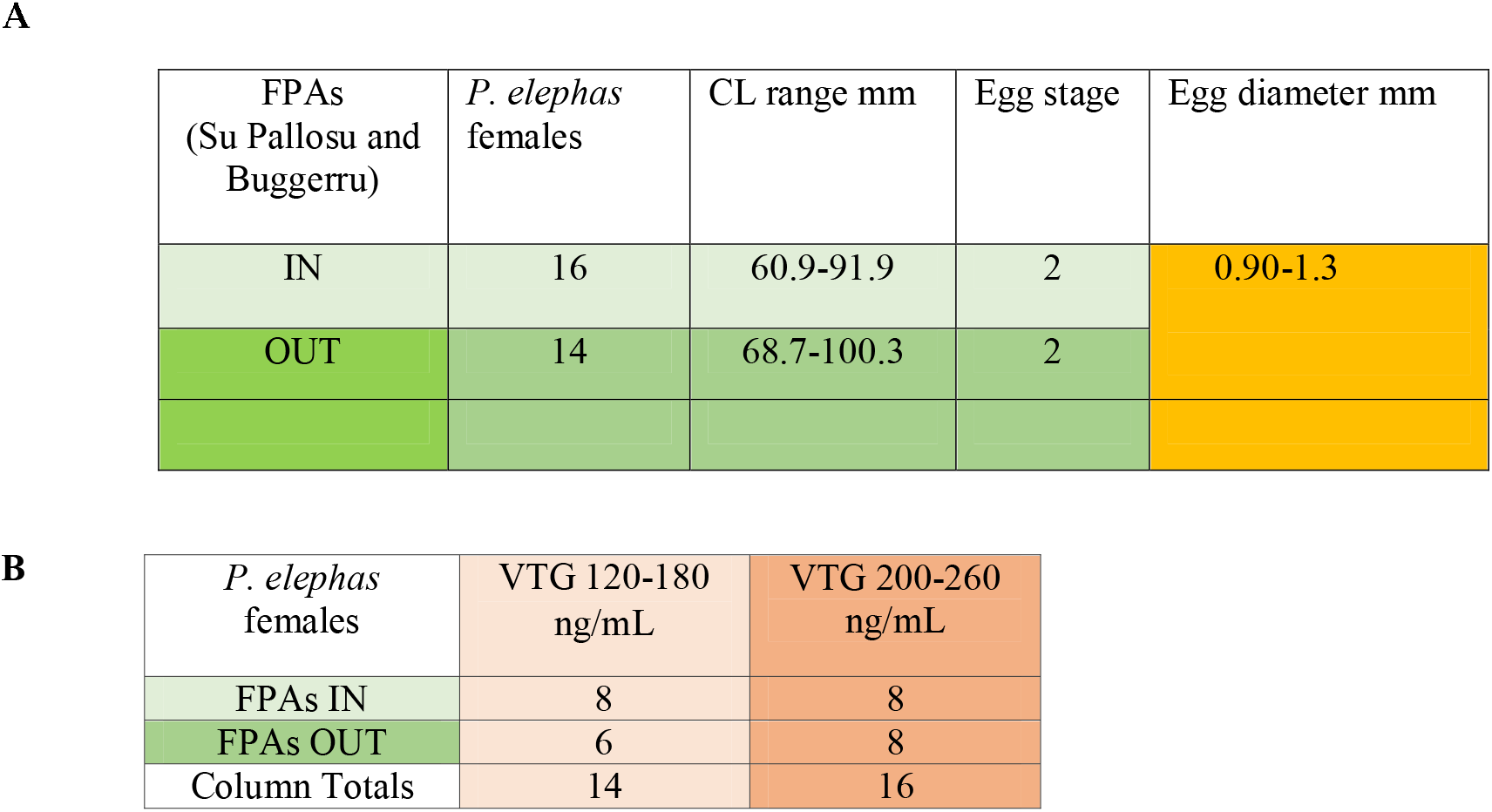
**(A)** Number of ovigerous P.elephas females captured inside (IN) and outside (OUT) FPAs, their size range and egg diameter. The presence of external eggs on the pleopods and carapace length (CL) was used as an indicator of maturity **(B)** Number of ovigerous females and the estimated VTG concentration range inside and outside FPAs. The VTG concentration in relationship with the inside and outside FPAs did not show statistic differences (Fisher’s exact test statistic = 1; p > 0.05).

## Notes

### Competing Interest Statement

The authors have declared no competing interest.

